# Temporal evolution of immunity distributions in a population with waning and boosting

**DOI:** 10.1101/253476

**Authors:** M. V. Barbarossa, M. Polner, G. Röst

**Affiliations:** Institute for Applied Mathematics, Heidelberg University, D-69120 Heidelberg„ Im Neuenheimer Feld 205, Germany; Bolyai Institute, University of Szeged, H-6720 Szeged, Aradi vértanúk tere 1, Hungary; Mathematical Institute, University of Oxford, Woodstock Road, OX2 6GG Oxford, United Kingdom

**Keywords:** Immuno-epidemiology, Waning immunity, Immune status, Boosting, Physiological structure, Delay equations, Basic Reproduction Number

## Abstract

We investigate the temporal evolution of the distribution of immunities in a population, which is determined by various epidemiological, immunological and demographical phenomena: after a disease outbreak, recovered individuals constitute a large immune population, however their immunity is waning in the long term and they may become susceptible again. Meanwhile, their immunity can be boosted by repeated exposure to the pathogen, which is linked to the density of infected individuals present in the population. This prolongs the length of their immunity.

We consider a mathematical model formulated as a coupled system of ordinary and partial differential equations, that connects all these processes, and systematically compare a number of boosting assumptions proposed in the literature, showing that different boosting mechanisms lead to very different stationary distributions of the immunity at the endemic steady state. In the situation of periodic disease outbreaks, the waveforms of immunity distributions are studied and visualized. Our results show that there is a possibility to infer the boosting mechanism from the population level immune-dynamics.

**AMS Classification:** 92D30, 34K60, 34K34, 37M05

## 1 Introduction

The outcome of an infection within an individual host depends on the specific pathogen and the status of the immune system of the host. At a larger scale, the outcome of an epidemics in a population is influenced by the ensemble of individual immunities. There are a number of processes in play that determine how these immunities change in time. Upon recovery from infection, some immune memory remains, which may persist for long time after pathogen clearance. Eventually, memory cells slowly decay, and in the long run recovered hosts could lose pathogen-specific immunity [20]. Waning immunity is possibly one of the contributing factors which cause, in particular in highly developed regions, recurrent outbreaks of infectious diseases such as chickenpox and pertussis. Immune memory can be boosted due to repeated exposure to the pathogen thus prolonging the time during which immune hosts are protected. Our goal in this paper is to monitor the distributions of immune memories in a population and track their temporal evolution, which provides very important insights about the interplay of individual and population level disease dynamics.

In the *SIR* framework, a population of hosts is divided into susceptibles (*S*), infectives (*I*) and recovered (*R*), and interactions among individuals from the different compartments are considered. Susceptibles are those hosts who either have not contracted the disease in the past or have lost immunity against the disease-causing pathogen. When a susceptible host gets in contact with an infective, the pathogen can be transmitted and the susceptible host may become infective. After the loss of immunity, an individual from compartment *R* transits back to the susceptible compartment, hence when waning of immunity is included, the model is called *SIRS*.

To account for immune system boosting, we structure the immunes according to their level of immunity. The high complexity of the immune status of an individual is simplified into a single parameter that reflects the strength of immunity in the sense that it indirectly indicates the duration of immunity until waning. The challenge due to secondary or multiple exposures to the pathogen initiates an immune response that results in a higher level of immunity.

The details are yet unclear how exactly the immune response and in particular a boost of the immune system work [20], most likely there is a range of mechanisms underlying these processes that are specific to each host and pathogen [1]. Laboratory analysis on vaccines tested on animals or humans suggest that the boosting efficacy might depend on several factors, among which the current immune status of the recovered host and the amount of pathogen he receives [1, 13]. In previous mathematical models it was assumed that a boost restores the maximal immune status, the same as after natural infection [2]. Few authors have assumed that at each new contact with a known pathogen, the immune system is boosted a little, with a small increase in memory cells [11]. Others have considered a combination of both possibilities [10].

To investigate the temporal evolution of how the distribution of these immunity levels change in the population, we propose a mathematical modeling framework along the lines of our previous works [4, 3]. The core of the model is a hybrid system of equations of SIRS type, in which the immune population is structured by the level of immunity, whereas the susceptible and the infective populations are non-structured. In [4] we investigated the well-posedness of the general model and its basic qualitative properties, whereas in [3] we considered a special case of the hybrid system in form of delay differential equations (DDEs) with constant and distributed delay. Here we focus on the immune response identifying several possible scenarios for immune system boosts.

We systematically compare a number of assumptions on the boosting mechanism that has been used in the literature. The goal is to observe the effects of different immune responses not only at an individual but also at population level, such as the stationary distribution of immunities in the case of an endemic disease, and the periodic change of these distributions in case of repeated disease outbreaks. To the best of our knowledge, there are no further studies about the temporal evolution of the distribution of immune statuses in a population, and the only paper that has predicted a specific immunity distribution from a mathematical model is [10].

In Section 2 we provide the details of the mathematical model, and present the numerical methods that are used for its solution. Four scenarios for immune boosting mechanisms are described and compared in Section 3, together with the parametrizations that lead either to a stable endemic state or oscillatory disease outbreaks. A careful and systematic numerical analysis is performed to understand the effect of various boosting mechanisms, and the results are summarized and discussed in Section 4.

## 2 Materials and Methods

### 2.1 The mathematical model

Let *S*(*t*) and *I*(*t*) denote the total population of susceptibles, respectively infectives, at time *t*. The total population shall be assumed to be in balance (hence normalized) *N*(*t*) ≡ 1, with birth rate equal to the natural death rate *d* ≥ 0 and no disease-induced death. We assume that newborns are all susceptible.

Contact with infectives (at rate *βI*) induces susceptible hosts to become infective themselves. Infected hosts recover at rate *γ >* 0, that is, 1*/γ* is the average infection duration. Once recovered from the infection, individuals become immune, however there is no guarantee for life-long protection. Immune hosts who experience immunity loss become susceptible again.

Let *r*(*t, z*) denote the density of immune individuals at time *t* with immunity level 
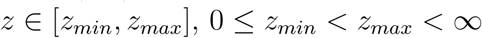
. The total population of immune hosts is given by

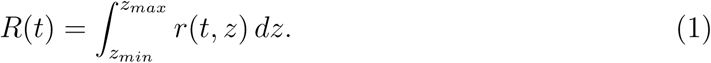

The parameter *z* describes the immune status and can be related to the number of specific immune cells of the host. The value *z_max_* corresponds to maximal immunity, whereas *z_min_* corresponds to the lowest level of immunity for hosts in the *R* compartment. We assume that individuals who recover at time *t* enter the immune compartment with maximal level of immunity *z_max_*. The level of immunity tends to decay in time and when it reaches the lower threshold *z_min_*, the host becomes susceptible again. However, contact with infectives, or equivalently, exposure to the pathogen, can boost the immune system from *z* ∈ [*z_min_*, *z_max_*] to any higher immune status, see Fig. 1.

**Figure 1:**
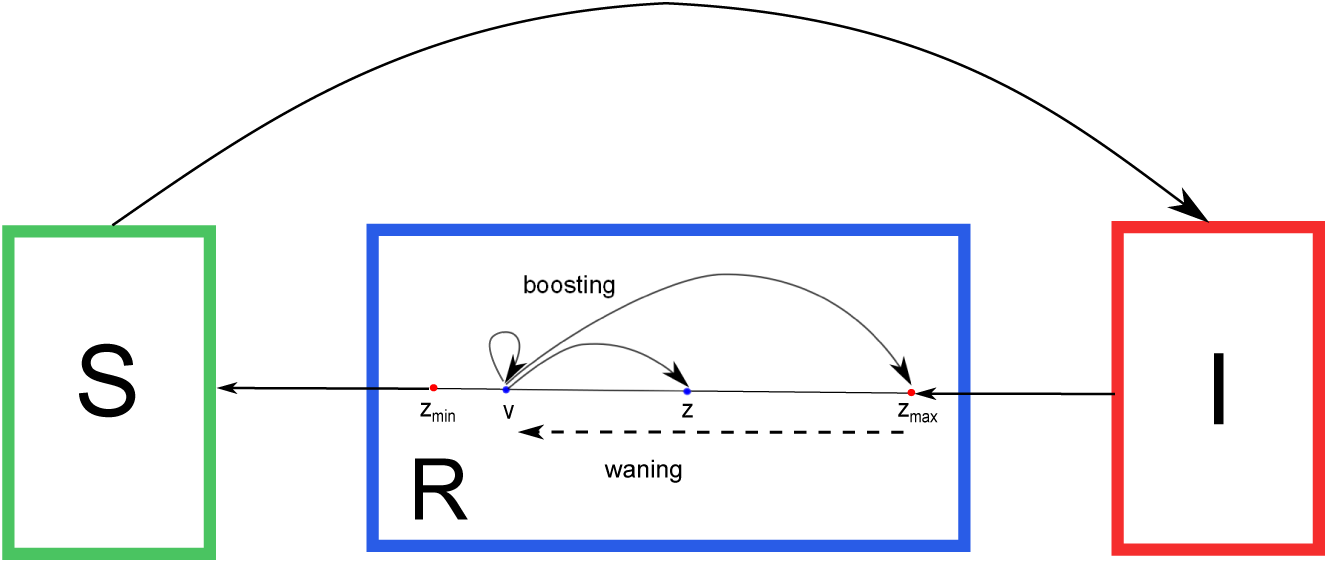
Sketch of the mathematical model (3)–(5). Susceptible hosts become infective after pathogen transmission. Infected hosts who recover enter the immune compartment *R* which is structured by the level of immunity. Natural infection induces the maximal level of immunity *z_max_*. Immunity decays in time and when the immune status reaches the minimal value *z_min_*, the recovered host becomes susceptible again. Meanwhile exposure to the pathogen can boost the immune system and prolong protection.

Given a host with *initial* immune status *v* ∈ [*z_min_*, *zmax*], let us denote by *Z^v^* ∈ [*z_min_*, *z_max_*] the *updated* immune status, which is achieved after new contact with the pathogen. The updated immune status *Z^v^* is modeled as a random variable taking values in 
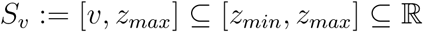
 with probability density function 
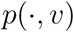
. The value *p*(*z*, *v*) represents the relative likelihood that a boost from immune level *v* evel *z* occurs. Secondary exposures to the pathogen might have no effects on the host’s immune system or might restore the immunity level induced by the disease (*z_max_*). In order to capture these particular aspects, in Section 3.1 we shall also consider limit cases in which the probability density function is a Dirac measure centered either on the current immune status or on the maximal level *z_max_*.

The immunity level decays in time at rate *g*(*z*), with *g* positive, smooth and bounded, which is the same for all immune individuals with immunity level *z*. In the absence of immune system boosting, an infected host who recovered at time *t*_0_ becomes again susceptible at time *t*_0_ + *τ*, where

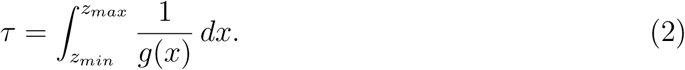

With the above assumptions, we obtain for *t >* 0 the following system of equations

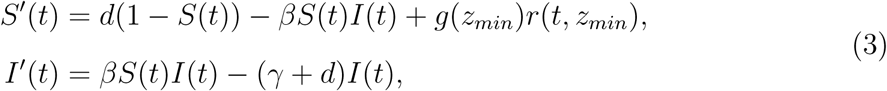

with initial values *S*(0) = *S*^0^ > 0, *I*(0) = *I*^0^ ≥ 0, coupled with a partial differential equation (PDE) for the immune population,

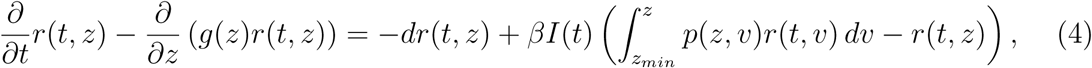

for *z* ∈ [*z_min_*, *z_max_*), with boundary condition

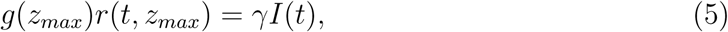

and initial distribution 
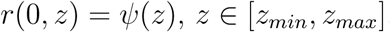
. A sketch of the full model is given in Fig. 1, whereas some possible boosting mechanisms are depicted in Fig. 2, and they will be discussed later in detail. The formal derivation of a slightly more general version of model (3)–(5) with variable total population size and disease-induced death is given in [4].

**Figure 2:**
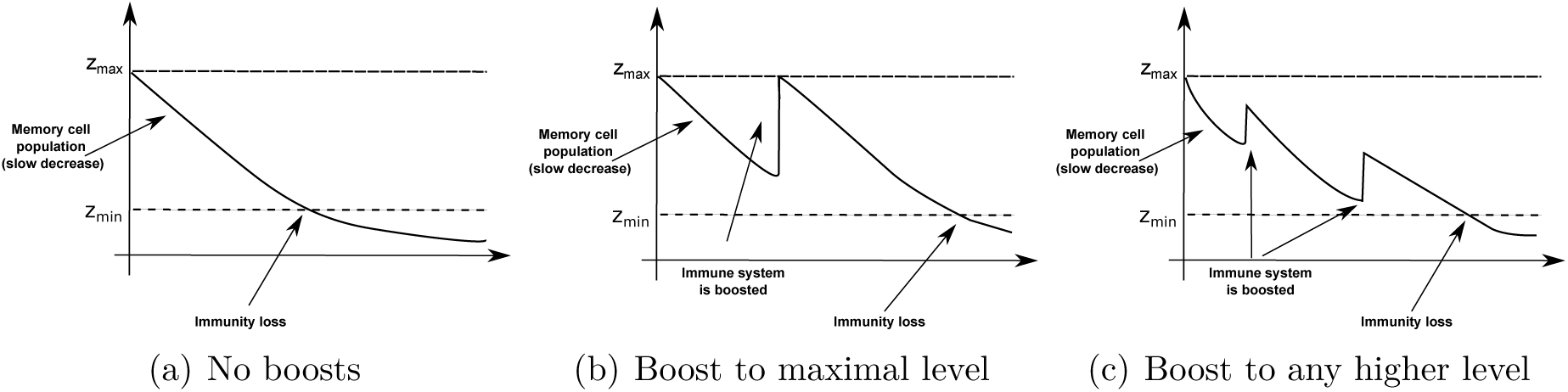
Exposure to the pathogen has a boosting effect on the immune system, whereby several possible scenarios are possible: (a) No boosting events (short: NOboost), the host who recovered at time t becomes susceptible at time t+τ, with τ > 0 given in equation (2). (b) Boosts restore disease-induced immunity, that is, the immune system is always boosted to the maximal level of immunity (short: MAXboost). (c) Variable boost, to any higher immune level.

### 2.2 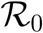

Before presenting further results we introduce the basic reproduction number 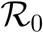 of model (3)–(5),

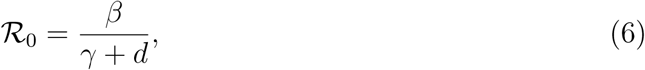

which indicates the average number of secondary infections generated in a fully susceptible population by one infected host over the course of his infection. The basic reproduction number is a reference parameter in mathematical epidemiology used to understand if, and in which proportion, the disease will spread among the population.

### 2.3 Numerical solution of the hybrid system

In the following, we outline the numerical method used to solve the hybrid system (3)–(5).

Consider (3)–(5) as a one dimensional, first-order nonlinear PDE system. In this section we refer to the independent variables (*t*, *z*) as time and space, respectively. As *S* and *I* do not depend on *z*, but only on time, the initial conditions are

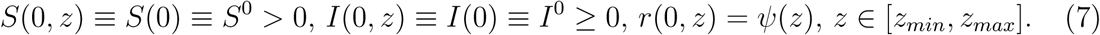

Since *g*(*z*) > 0 for all *z* ∈ [*z_min_*, *z_max_*], the boundary condition for *r* is imposed in (5) at the inflow boundary, i.e., at *z* = *z_max_*. All together we have an initial-boundary value problem (IBVP). For the numerical integration of the IBVP we employ the MATLABⓇ code *hpde* [16], developed to solve IBVPs for first order systems of hyperbolic PDEs in one space variable and time. The *hpde* routine implements Richtmyer’s two-step variant of the Lax-Wendroff method [12, 15], which is well established to solve hyperbolic PDEs. This scheme is explicit and second order accurate in both space and time. To compute the numerical solution of our IBVP, we have modified the code *hpde* so that the integral term in (4) is efficiently implemented, preserving the second order accuracy of the Lax-Wendroff scheme. In Section 3.3 we have verified the spatial order of accuracy for a number of computations.

In the computation of the numerical solutions of the IBVP, we used an equidistant mesh

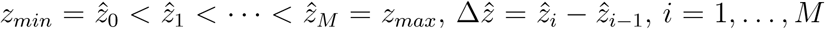. The initial function on this mesh is given by 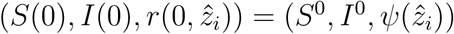.

For an explicit scheme for a hyperbolic systems, a necessary condition for stability is the Courant-Friedrichs-Lewy condition, which for our system means

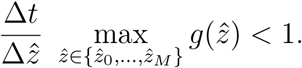

We define the *boosting matrix* Π as the numerical discretization of the probability density function *p* on the equidistant mesh 
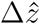
. Each entry *π_ij_* = Π(*i*, *j*) of the boosting matrix represents the probability that 
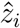
 is the updated immune level, given initial immune level
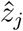
 that is, 
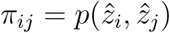
. It follows that *π*_*ij*
_ ∈ [0, 1], with *π*_*ij*_ = 0 if *j* > *i* and

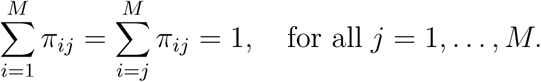

## 3 Results and Discussion

### 3.1 Scenarios for the immune response

Let us consider an immune host who has immune status *v* at the moment of re-exposure to the pathogen. We shall investigate the following possible scenarios for immune response in case of secondary exposure to the pathogen.

#### NOboost

Assume that the immune system does not respond to reinfection, that is, we observe only waning of immunity after recovery, Fig. 2(a). This corresponds to the limit case in which the boosting probability function is simply a Dirac measure with support on the initial immune level. It follows that equation (4) reduces to

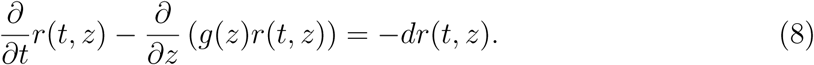

Recall the definition of *τ* > 0 in (2). The transport equation (8) with boundary condition (5) is solved along characteristics and we obtain

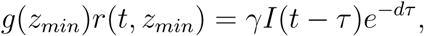

meaning that *τ* time after recovery immune hosts who did not die become susceptible again. In turn, we find a delay term in the equation for *S* and have a classical SIRS model with constant delay (cf. [17])

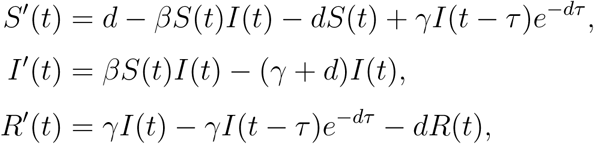

where *R*(*t*) is the total immune population at time *t* as in (1).

#### MAXboost

Assume that at any new encounter with the pathogen the immune system of a recovered host is boosted in such a way that the disease-induced (maximal) immunity is restored, Fig. 2(b). This corresponds to the limit case in which the boosting probability function is a Dirac measure with support on the maximal immune level (*z*_*max*_). As all boosted individuals are transferred to the boundary, the integral term in equation (4) moves to the boundary condition, yielding

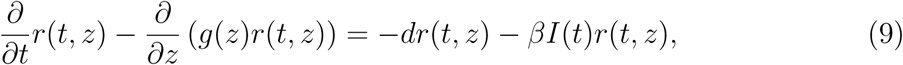

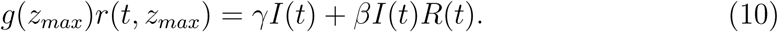

The BVP (9)–(10) can be solved along characteristics and one obtains for *t* ≥ 0 a system of delay equations with constant and distributed delay:

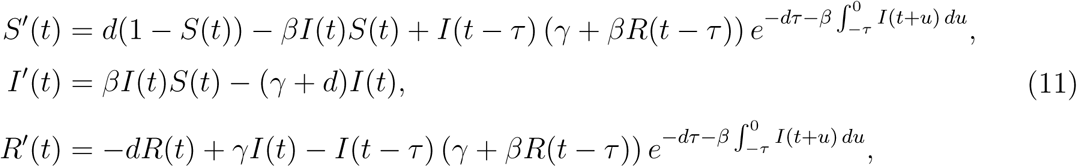

with *τ >* 0 as defined in (2), and with given initial functions 
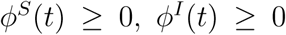
,
*ϕ^R^*(*t*) ≥ 0, such that 
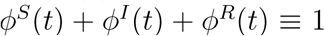
 for all *t* ∈ [−*τ,* 0]. The formal derivation of system (11)) is given in [3, 4].

#### ANYboost

We assume that the immune system is boosted to any better immunity level with uniform probability. That is, 
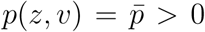
 with 
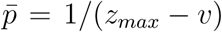
, for all *z* ∈ [*v*, *z_max_*]. The boosting matrix for the numerical implementation is shown in Fig. 3(a).

**Figure 3:**
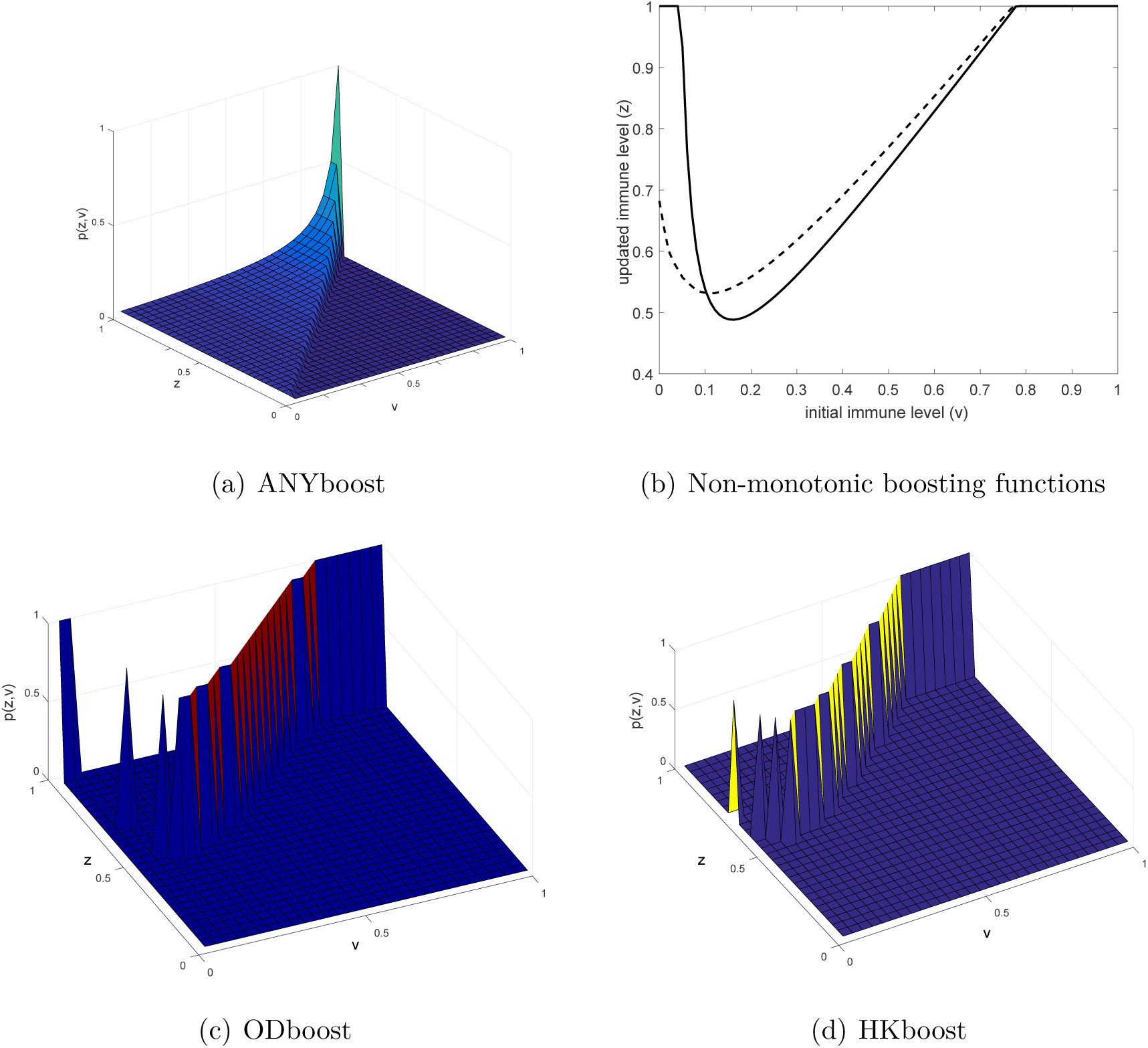
Non-monotonic immune response mechanisms. (a,c,d) Examples of different boosting matrices Π for the numerical implementation of *p*(*z, v*), the probability that individuals with immune level *v* at exposure are boosted to level *z* ≥ *v* after exposure. The immune status is represented in [*z_min_*, *z_max_*] = [0, 1] and discretized on an equidistant mesh with 30 points. (b) The dotted curve represents the boosting mechanism inspired by [10], the solid curve the one inspired by [8].

#### HKboost and ODboost

Previous works have proposed non-monotonic functions for modeling the immune response to secondary exposure. In [9, 10], a mathematical model for in-host dynamics during measles infection was proposed. The model explicitly considers the relation between the immune level at time of exposure and the level of memory cells after exposure. Heffernan and Keeling [9, 10] suggest that this relation is non-monotonic. Indeed, although the level of immunity after exposure is always greater than the level at the time of exposure, the boosted level starts high, decreases and then increases again. We shall denote by HKboost the boosting function from [9, 10]. For the numerical implementation of HKboost we first extract values from Fig. 2(d) in [10], which shows the relation between the initial and the updated immune status. Then we normalize the immune status interval ([*z_min_*, *z_max_*] = [0, 1]) and extend the boosting function to [0.75,1], assuming that for high initial immune status, the immune system is always boosted to the maximal level, see Fig. 3(b). In terms of our model coefficients, for any initial immunity *v*, the atomic measure of the updated immunity is concentrated on *f*(*v*), where *f* is the boosting function represented by the dotted curve in Fig. 3(b). The corresponding boosting matrix is shown in Fig. 3(d).

Immune boosting in pertussis was considered in [8], where a non-monotonic boosting function similar to the one in [9, 10] was proposed. The relation between the level of memory cells before (*v*) and after (*z*) re-exposure is governed by the function

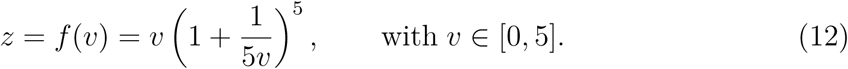

De Graaf et al. [8] suggest that a small jump from *v* to *z* corresponds to a mild infection that causes no harm but only boosts the antibody level of the individual, whereas a large jump corresponds to a severe infection that may also cause disease. We shall use the boosting function in (12) for simulations, previous normalization to [*z_min_*, *z_max_*] = [0, 1], and small modifications which allow to have boundedness. In detail, we assume that for very low initial immunity (0 ≤ *v <* 0.075) and for large initial immunity (0.8 < *v* ≤ 1), the immune status is always boosted to the maximal level *z* = *z*_*max*_ = 1, whereas for *v* ∈ [0.075, 0.8] the boosting probability is atomic along the graph of the boosting function *f* in (12), as above. The resulting boosting function is given by the solid curve in Fig. 3(b), and the corresponding boosting matrix is shown in Fig. 3(c).

The two mechanisms for immune response HKboost and ODboost are governed by rather similar boosting mechanisms and we shall see that the solutions behave accordingly.

### 3.2 Parameter values

For the numerical computations below, we set the birth rate and natural death rate *d* = 0.02, and the initial conditions *ψ*
_0_(*z*) = 0.05 for *z* ∈ [*z_min_*, *z_max_*] = [0, 1], *I*^0^ = 0.01 and *S*^0^ = 0.94. This means that we assume that 1% of the total population is infectious at the beginning of our observations, while 5% is immune to the pathogen and the level of immunity is equally distributed.

Concerning the disease dynamics, we set *γ* = 3 and *g*(*z*) = 0.5 for all *z* ∈ [*z_min_*, *z_max_*], corresponding to *τ* = 2 when the model can be reduced to DDE.

We allow the basic reproduction number 
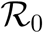
 as given in (6) to vary, in order to show both solutions that converge to an endemic equilibrium, and solutions that produce periodic oscillations.

### 3.3 Stable endemic equilibrium

When the basic reproduction number is sufficiently small, the solution converges to an endemic equilibrium for all the boosting mechanisms presented in Section 3.1. We compare in Fig. 5 the numerical solution corresponding to different boosting mechanisms. In Fig. 5(a) we show the component *I* of the solution, which indicates how the number of infective evolves in time and rapidly converges to an equilibrium. Changing the boosting mechanism has small effects on the infective population: numerical solutions show the same qualitative behavior, with endemic equilibria deviating less than 1% from each other. In particular, we see that the non-monotonic boosting mechanisms, ODboost, HKboost and ANYboost, yield quantitatively equivalent solutions for the infective population. The latter solution curves are bounded from below by the solution of MAXboost and from above by the solution of the NOboost problem.

In Fig. 5(b), for each boosting mechanism we visualize the stationary distribution 
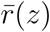
 corresponding to the endemic equilibrium in Fig. 5(a). This indicates how immunity is distributed among the *R*−population at the endemic equilibrium. It is visible that the choice of the boosting mechanism importantly affects the stationary distribution. To understand why this happens, let us consider the case of a constant immune decay rate, 
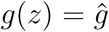
 for all *z* ∈ [*z_min_*, *z_max_*]. For certain choices of the probability density function *p* one can calculate the stationary distribution 
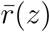
 explicitly. For example, in the absence of immune boosts (NOboost), we have

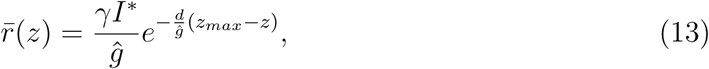

where *I*\* is an endemic equilibrium of the system (3), (8), with boundary condition (5).

In case of boosts to maximal immune level (MAXboost), from (9)–(10) we have

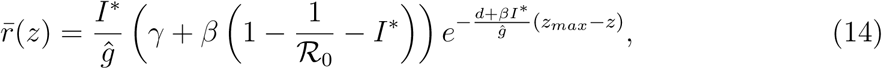

with the basic reproduction number 
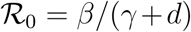
, as in (6). With the parameter values indicated in Section 3.2, and 
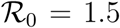
 the curves in (14) and (13) are almost straight lines, as it can be observed in Fig. 5(b).

We use the analytic stationary solution (14) of system (3)–(4), (9)–(10) to verify the order of ”spatial” accuracy (i.e., with respect to the variable *z*) of the numerical discretization. On different meshes, we compute numerically the stationary solution of the MAXboost system (3)–(4), (9)–(10). Although, we know the explicit solution for the stationary distribution 
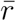
, there is no explicit analytical formulation for the endemic equilibrium *I*\*, which can only be determined numerically (see also [3]). We fix *I\** to the value computed on the finest mesh (*M* = 14000 equidistant mesh points) and we insert this value into the formula (14) of the stationary distribution. Then, we compute the maximum error between the analytical and the numerical solution on different mesh refinements and observe that our method preserves the second order accuracy of the Lax-Wendroff scheme, see Fig 4.

**Figure 4:**
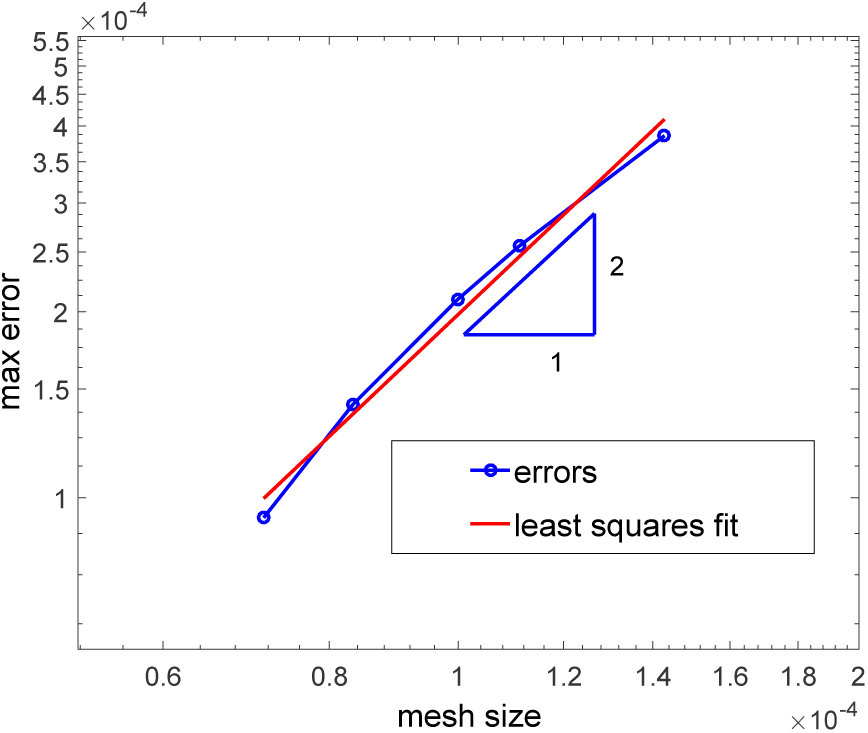
Spatial accuracy of the numerical discretization, with error represented in a log-log plot. The method is second order accurate.

### 3.4 Sustained oscillations, repeated outbreaks

When 
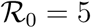
 and all other parameter values are as indicated in Sect. 3.2, the system (3)–(5) shows periodic oscillatory behavior, independent on the boosting mechanism. In Fig. 6 we compare the *I* component of the solution of the system (3)–(5) for five different immune responses. Further, for all kinds of immune response, we compare in Fig. 7 the distribution of immune status among the population for fixed times (*t* = 1, 5, 10, 20 years) after the beginning of the observation. Results illustrated in Fig. 6, Fig. 7 and Table 1 indicate that also in the case of periodic oscillations the solution of problems governed by ODboost and HKboost, are qualitatively and quantitatively equivalent (minor differences might be due to computational errors and we consider them negligible). Therefore, in the following, we show only results related to NOboost, ANYboost, MAXboost and ODboost, which characterize four different immune responses with qualitatively different results. Fig. 8 shows the solution *r*(*t, z*) with respect to time and immune status for the four selected boosting mechanisms.

**Figure 5:**
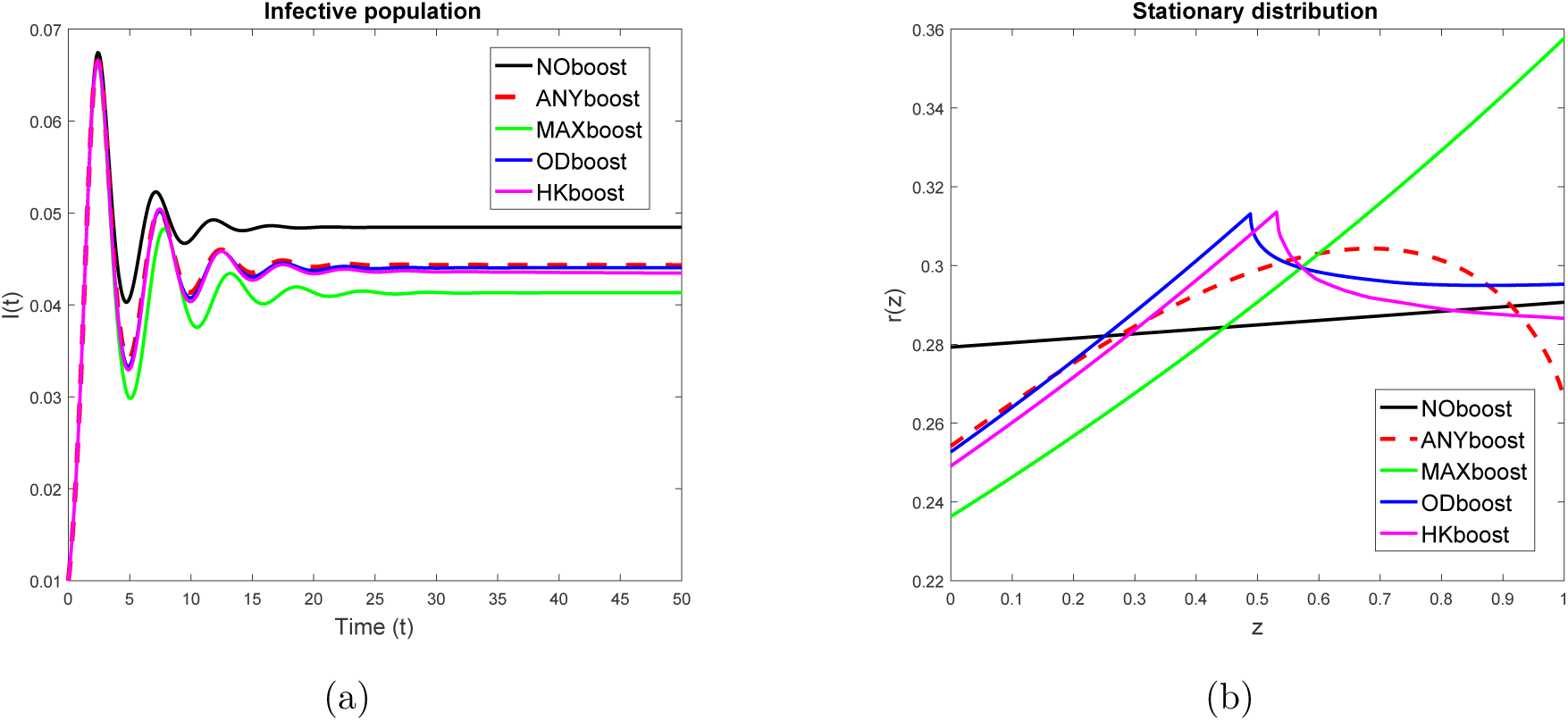
Comparison of different boosting mechanisms in case of a stable endemic equilibrium. (a) The infective population in case of NOboost (black curve), MAXboost(green), ANYboost(red dotted), ODboost (blue) and HKboost (magenta). (b) The stationary distribution in the immune population 
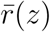
) corresponding to (a).

**Figure 6:**
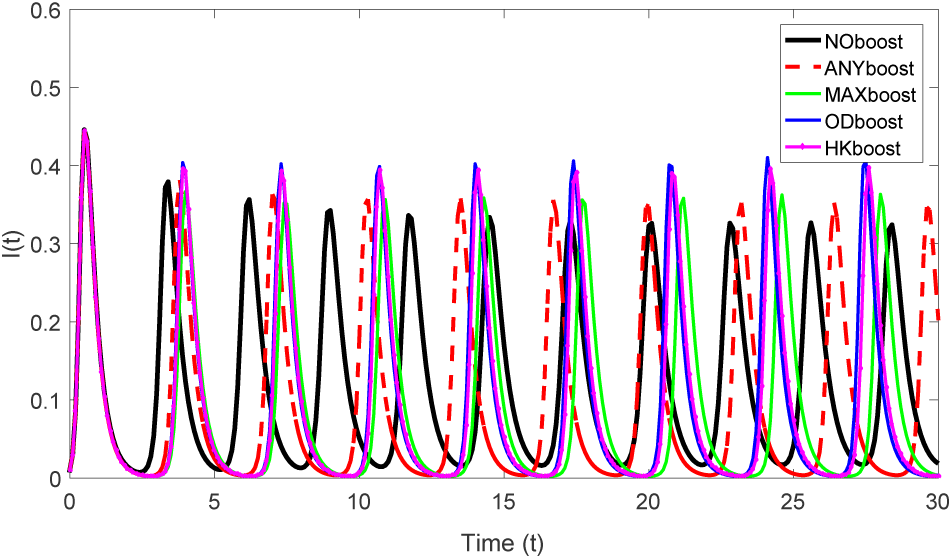
Infective population *I*(*t*) for different boosting mechanisms. Also in case of periodic oscillations it is evident that the solution of problems governed by the boosting mechanisms ODboost and HKboost are quantitatively equivalent.

**Figure 7:**
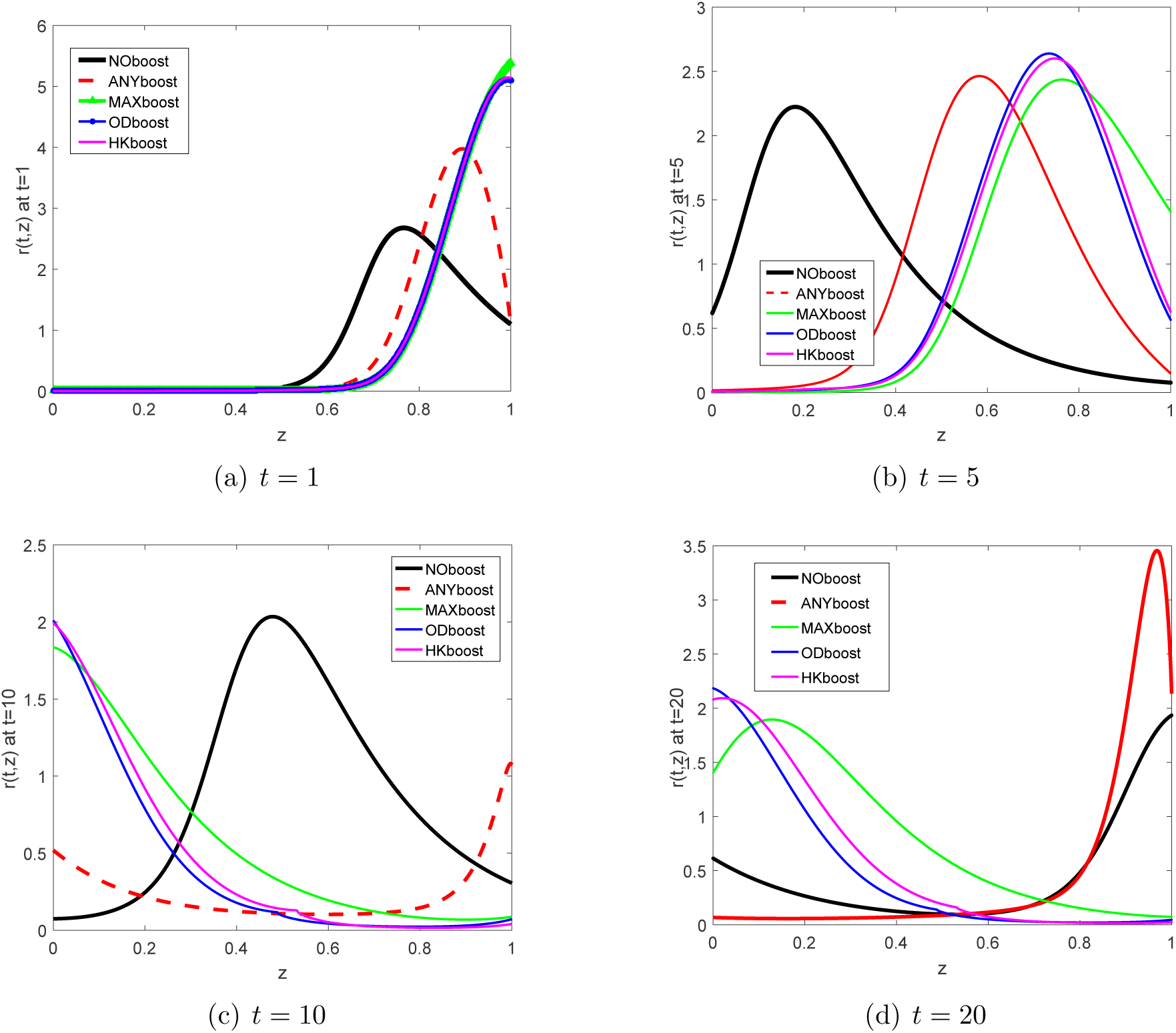
Comparison of the effects of different boosting mechanisms on the distribution of immunity *r*(*t*, z*) at fixed times *t\** = 1, 5, 10, 20 years after the beginning of the observations.

**Figure 8:**
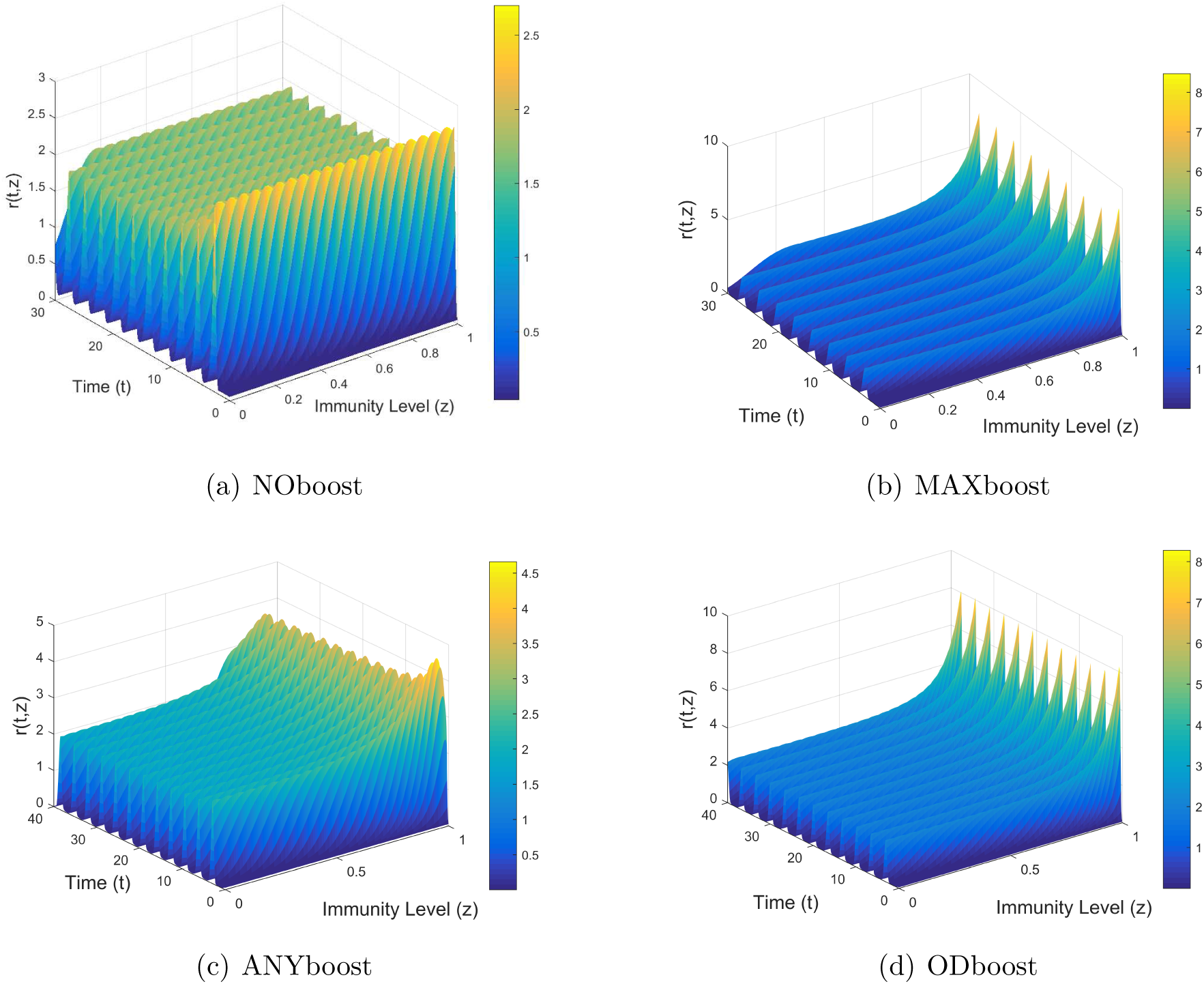
The *r*(*t*, *z*) solution for different boosting mechanisms. (a) Waning immunity, absence of immune boosts. (b) Boost to maximal level of immunity. (c) Uniform probability of boosting to any higher immunity level. (d) Non-linear boost according to Fig. 3(c).

**Table 1:** Quantitative comparison of the periodic oscillatory solutions (*I*-component) for changing immune boost mechanism. Oscillation amplitudes and periods, as well as lowest and highest incidence values are reported.

|  | Amplitude | Period | min(I) | max(I) |
| --- | --- | --- | --- | --- |
| **NOboost** | 0.3088 | 2.766 | 0.0169 | 0.3257 |
| **MAXboost** | 0.3588 | 3.4 | 0.0024 | 0.3612 |
| **ANYboost** | 0.3430 | 3.2 | 0.0036 | 0.3466 |
| **ODboost** | 0.4105 | 3.35 | 0.0023 | 0.4129 |
| **HKboost** | 0.3926 | 3.35 | 0.0024 | 0.3950 |

In Fig. 9 we visualize the contour plots of the *r* component of the solution for different immune response mechanisms. Entries of the solution matrix *r* are interpreted as heights with respect to the (*z*, *t*) plane, isolines are calculated and displayed using colors corresponding to the colormap on the right.

**Figure 9:**
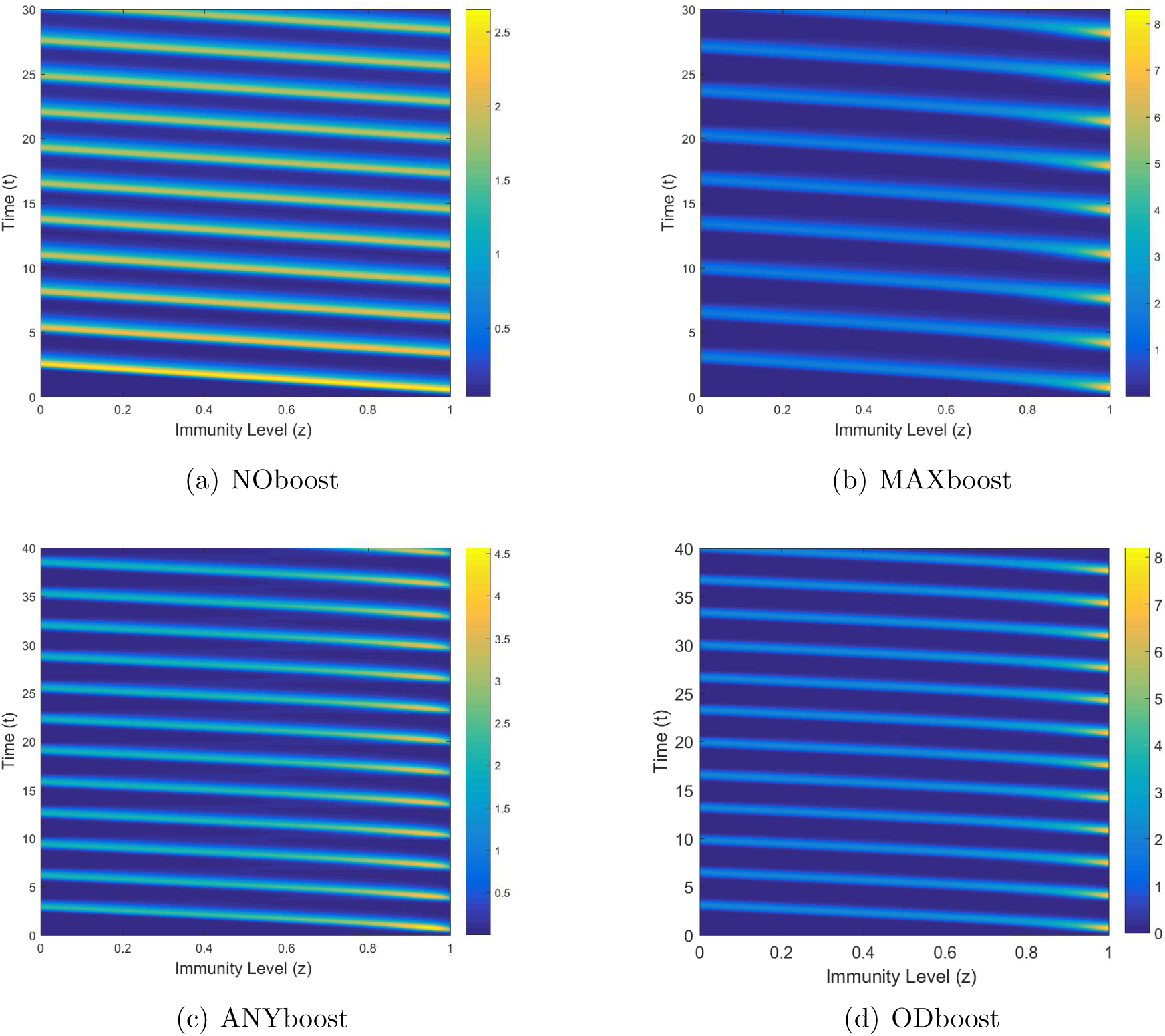
Contour plot of the *r* solution for different boosting mechanisms. (a) Waning immunity, no boosts. (b) Boost to maximal level of immunity. (c) Uniform probability of boosting to any higher immunity level. (d) Non-linear boosting function as in Fig. 3(c).

In Fig. 10 we compare the immune distribution *r*(*t*, *z*) corresponding to different levels of an outbreak, for different boosting mechanisms. Fig. 10 (a) shows four points on a typical infective curve (*I*) which we shall consider for comparison. Point A corresponds to the infection peak, that is the time point at which the infective population is at its maximal value. Point B and point D correspond to intermediate levels in the infective population, just after, respectively, just before an outbreak peak. Point C corresponds to the time point at which the infective population is at its minimal level. We see in Fig. 10(b–e) that the different immune responses yield qualitatively similar immune distributions with immunity waves moving from right to left. At the infective peak (Fig. 10(b)), following primary infection and re-exposure to the pathogen, most of the population has a very high immune status (*z* ∈ [0.8, 1]) which slowly decays, reaching the level at which most of the population has a low to intermediate immune level (*z* ∈ [0, 0.6]), Fig. 10(d). Just before a new outbreak, most of the hosts have very low (*z* ∈ [0, 0.2]) or very high (*z* ∈ [0.8, 1]) immunity, Fig. 10(e).

**Figure 10:**
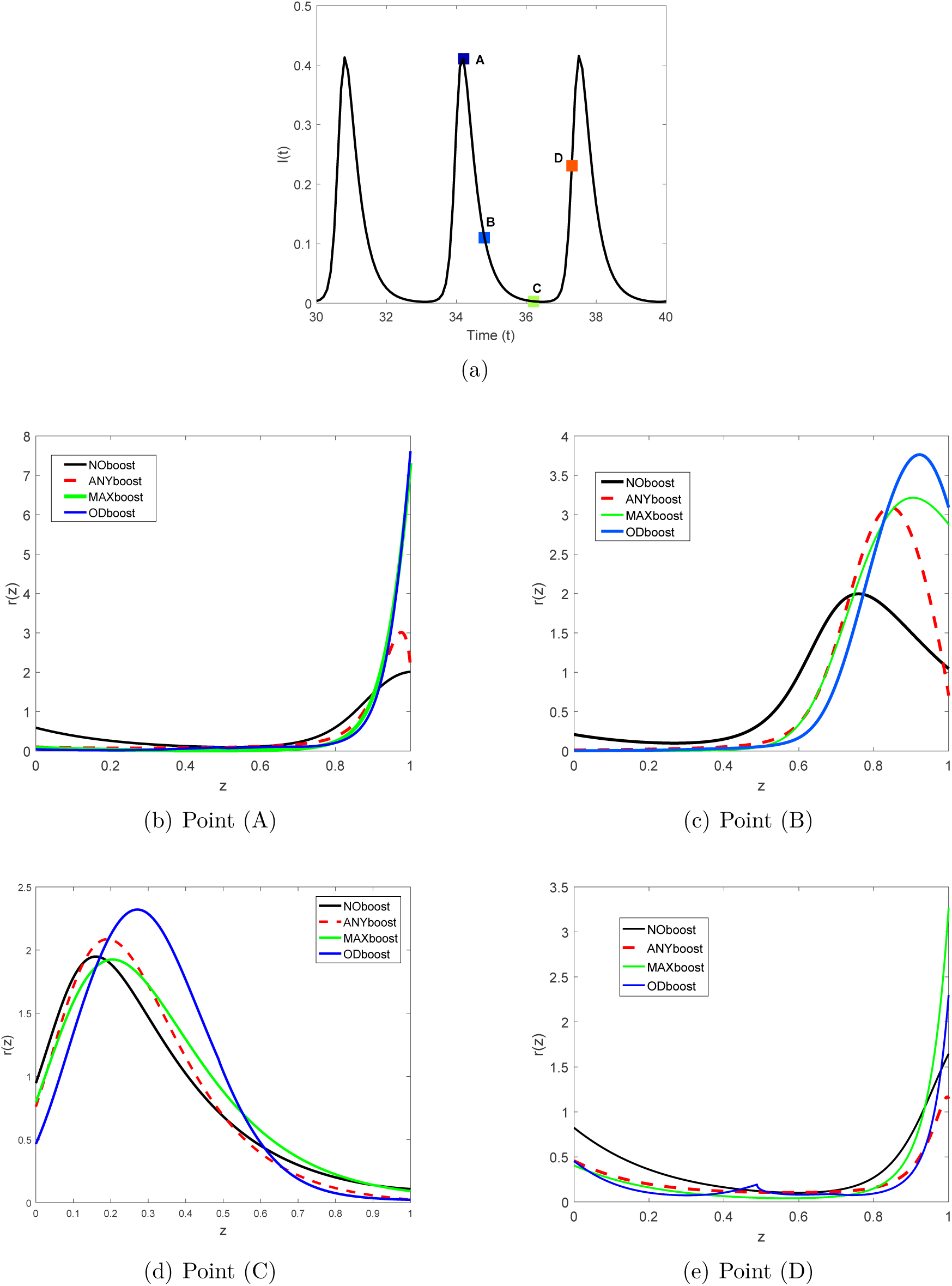
Comparison of immune distribution for different boosting mechanisms at different levels of an outbreak. (a) The *I* solution and the four reference points to be considered for comparison: (A) the infection peak, (B) an intermediate level after the outbreak, (C) the minimal infection level, (D) just before a new outbreak. (b-e) Immune distribution corresponding to the points (A)-(D).

## 4 Conclusion

Understanding the role of immune system boosting due to the interplay of in-host and between hosts dynamics is a central point in immuno-epidemiology of infectious diseases. Employing the mathematical model (3)–(5), in this work we have systematically investigated the effects of different (in-host) immune responses at population level. The boosting mechanisms studied here are in part taken or inspired by previous literature, e.g. on measles [9, 10], pertussis [8] and influenza [21].

Our results indicate that the in-host immune response, i.e. the individual boosting mechanism, has important effects on the qualitative dynamics of the epidemics. The numerical results show that observing the distribution of immune level among recovered individuals, one can reconstruct the underlying boosting mechanism. When the trajectories of the system (3)–(5) approach an endemic equilibrium (Fig. 5(b)), the distribution of immunity uniquely identifies the boosting mechanism governing the immune response in case of secondary infections. Also for diseases with repeated outbreaks it is possible to infer the immune boosting mechanism from the temporal evolution of immunity, though in this case it is convenient to look at the dynamic of the immune population over time (Fig. 8, Fig. 9), rather than at the immune distribution (Fig. 10).

This paper presents a novel mathematical study of temporal evolution of immune status in a population. Despite the large number of publications in mathematical epidemiology or mathematical immunology, to the best of our knowledge, there have been only few authors who have effectively combined in-host dynamics and processes at population level [9, 10, 19, 14], though only in [9, 10] the temporal evolution of immunity has been considered. If we compare the stationary distribution of immunity of HKboost in Fig. 5(b) with Fig. 3(c) in [10], we observe that the results are rather different, the curve in [10] being smoother. We conjecture that the differences are on the one hand due to the choice of the parameters in the system, on the other hand to the different nature of the mathematical models. In [10], a large system of ordinary differential equation is presented, including exposed hosts and immune structure in all compartments, not only in the recovered ones as it is in our work. Here, we focused on a simple model for susceptible-infective-recovered dynamics in order to understand the role of immune boosting and waning immunity. We plan to extend the model (3)–(5), by means of physiologically structured susceptible and exposed populations.

Parameter values used for all numerical simulations presented in this work are not inspired by a specific infectious disease, as we purely focused on the effects of waning and boosting. Nevertheless, the model is suitable to describe several infectious diseases, prior proper choice of the parameters. Certain studies, e.g. [11, 7], introduced a further parameter *κ* to express a relative force of infection for reexposure, so they represent the rate of boosting by the term (*κβI*). This is reasonable if we think that during contacts, boosting occurs with a different chance compared to primary infection. In this paper we have assumed that the force of infection in secondary exposure is the same as in primary exposure (*κ* = 1).

The in-host dynamics described in this paper is rather coarse. The immune status of recovered individuals is represented by a single scalar quantity (*z*) which does not specifically represent any kind of cells of the immune system. We are aware of the fact that the immune system is indeed very complex, and careful mathematical modeling should also take into consideration nonlinear in-host processes and interactions among the different players of the innate and adaptive immune system. Nevertheless, such a fine description would quickly lead to a complicated mathematical model, making it hard to achieve any reliable analytical or numerical result at population level.

A further limitation of the model is the assumption on the sharp threshold *z_min_*, which defines the criterion for transition from immune to susceptible compartment. It is plausible that this transition does not occur in such an on-off manner, but it is rather a continuous process; that is, recovered individuals with a certain critically low level of immunity are less likely to get immune boost and more likely to experience a new infection, as suggested in [8]. Moreover, it is known that the immune system is differently reactive at different life stages, in particular immune boosts are weaker in children and elderly than in adults (see e.g. the case of pertussis considered in [18]). We plan to extend the model including age heterogeneity.

Immune boosts occur not only after natural infection but also in case of vaccine-induced immunity. Current vaccination schedules include ”boosters” whose goal is the prolongation of protection against a certain pathogen [6]. The mathematical model (3)–(5) can be extended to include vaccination and natural boosts induced in vaccinated hosts by contact with infectives, see [5]. Similar investigation as proposed in this paper can be repeated in the context of vaccination.

Our results indicate that immune system boosts from natural secondary infections importantly affects the immune dynamics at population level. We hope that our findings will stimulate future research in controlling infectious diseases via vaccination and vaccine design.

## Acknowledgements

MVB is supported by the European Social Fund and by the Ministry of Science, Research and Arts Baden-Württemberg. The work of MP was supported by the János Bolyai Research Scholarship of the Hungarian Academy of Sciences, the EU-funded Hungarian grant EFOP-3.6.2-16-2017-00015 and NKFI KH 125628. GR was supported by Marie Sklodowska-Curie Grant No. 748193.

